# Identifying protein subsets and features responsible for improved drug repurposing accuracies using the CANDO platform

**DOI:** 10.1101/405837

**Authors:** William Mangione, Ram Samudrala

## Abstract

Drug repurposing is a valuable tool for combating the slowing rates of novel therapeutic discovery. The Computational Analysis of Novel Drug Opportunities (CANDO) platform performs shotgun repurposing of 3,733 drugs/compounds that map to 2,030 indications/diseases by predicting their interactions with 46,784 protein structures and relating them via proteomic interaction signatures. The accuracy of the CANDO platform is evaluated using our benchmarking protocol that assesses indication accuracies based on whether or not pairs of drugs associated with the same indication can be captured within a certain cutoff, which is a measure of the drug repurposing recovery rate. To identify subsets of proteins that exhibit the same therapeutic effectiveness as the full set, groups of 8 proteins were randomly selected and subsequently benchmarked 50 times. The resulting protein sets were ranked according to average indication accuracy, pairwise accuracy, and coverage (count of indications with non-zero accuracy). The best 50 subsets of 8 according to each metric were progressively combined into supersets after each iteration and benchmarked. These supersets yield up to 14% improvement in benchmarking accuracy, and represent a 100-1,000 fold reduction in the number of proteins relative to the full set. Protein supersets optimized using independent compound libraries derived from the full library were cross-tested and were shown to reproduce the performance relative to using all 46,784 proteins, indicating that these reduced size supersets are broadly applicable for characterizing drug behavior. Further analysis revealed that sets comprised of proteins with more equitably diverse ligand interactions are important for describing drug behavior. Our work elucidates the role of particular protein subsets and corresponding ligand interactions that play a role in computational drug repurposing, and paves the way for the use of machine learning approaches to further improve the accuracy of the CANDO platform and its repurposing potential.

**Author summary:** Drug repurposing is a valuable approach for ameliorating the current problems plaguing drug discovery. We introduce a novel protein subset analysis pipeline that allows us to elucidate features important for drug repurposing accuracies using the Computational Analysis of Novel Drug Opportunities (CANDO) platform. Our platform relates drugs based on the similarity of their interactions with a diverse library of proteins. We subjected all proteins in the platform to a splitting and ranking protocol that ranked protein subsets based on their benchmarking performance. Further analysis of the best performing protein subsets revealed that the most useful proteins for describing how small molecule compounds behave in biological systems are those that are predicted to interact with a structurally diverse range of ligands. We hypothesize that this is a consequence of the multitarget nature of drugs and, conversely, the implied promiscuity of proteins in biological systems. These results may be used to make drug discovery more accurate and efficient by alleviating some of its bottlenecks, bringing us one step further in better understanding how drugs behave in the context of their environments.

## Introduction

Common strategies in drug discovery include forward pharmacology [1] and rational drug design [2]. In the former, a library of compounds is screened, typically in high throughout manner, for certain phenotypic effects *in vitro*. In the latter, compounds are virtually screened against a predetermined biological target and high confidence hits are then assayed for a desired modulation. In both cases, the hits obtained are then assessed for effectiveness *in vivo* and proceed to clinical trials for eventual FDA approval if successful at each stage. This iterative process can cost billions of dollars and up to 15 years per drug [3]. These approaches do not consider the promiscuity of approved drugs in the context of indications/diseases within living systems (evidenced by side effects present for all small molecule therapies [4, 5]), dooming many novel therapeutics to fail. With the second-leading cause of putative drug attrition being adverse reactions [6], there is great utility in finding new uses for already approved drugs, which is formally known as drug repurposing or repositioning [7, 8].

We have developed the Computational Analysis of Novel Drug Opportunities (CANDO) platform [9–11] to address these drug discovery challenges. One fundamental tenet of CANDO is that drugs interact with many different proteins and pathways to rectify disease states, and this promiscuous nature is exploited to relate drugs based on their proteomic signatures [9, 12–15]. These signatures are typically determined via virtual molecular docking simulations that are applied to predict compound-protein interactions on a proteomic scale. Using a knowledge base of known drug-indication approvals/associations, we can identify putative drug repurposing candidates for a particular indication based on the similarity of their proteomic interaction signatures to all other drugs approved for (or associated with) that indication. When a particular indication does not have any approved drug, the library of human use compounds present in CANDO is screened against the tertiary structures of all relevant and tractable proteins obtained by x-ray diffraction or homology modeling from a particular organismal proteome to suggest new treatments that maximize binding to the disease-causing proteins and minimize off-target effects. High-confidence putative drug candidates generated by CANDO using both approaches have been prospectively validated preclinically for a variety of indications, including dengue, dental caries, diabetes, hepatitis B, herpes, lupus, malaria, and tuberculosis, with 58/163 candidates yielding comparable or better therapeutic activity than standard treatments [12, 13, 16, 17].

To date, putative drug candidates generated by CANDO have been based on simple comparison metrics, primarily the root mean square deviation (RMSD) of the binding scores present in a pair of drug-proteome interaction signatures. Our platform is evaluated using a benchmarking method that assesses per indication accuracies based on whether or not other drugs associated with the same indication can be captured within a certain cutoff in terms of similarity to a particular drug approved for that indication. Incorporating machine learning, which is continuing to prove its utility in many aspects of biomedicine [18–20] including drug discovery and repurposing [21, 22], into the CANDO platform to increase benchmarking accuracies and therefore its predictive power is of importance. Various algorithms can be incorporated (for example, neural networks, support vector machines, and decision trees), but the well documented issues described by the curse of dimensionality [23, 24] will plague any choice in the current (v1) implementation of CANDO, especially considering the extremely large number of features (≈50,000 proteins) within each compound-proteome interaction signature vector. Given the relatively few training samples (an average of ≈9 drugs per indication), a machine learning approach to train how drugs interact with proteomes is a much easier task with a vastly reduced set of proteins.

Therefore, in this study we set out to find a reduced set of proteins that can therapeutically characterize compounds as well as using the full protein library, with the eventual goal of utilizing them in future machine learning experiments. Our strategy involves using a brute force or greedy feature selection, where the full protein library was split into smaller subsets that were subsequently benchmarked and ranked according to their performance by different metrics. The best performing subsets were recombined into supersets and benchmarked again, which produced substantial improvement with all the benchmarking metrics. Further analysis of the best performing protein subsets and supersets revealed that those that contained proteins predicted to bind to a more equitably diverse ligand structure distribution were strongly associated with increased benchmarking performance, indicating that drugs approved for human use have a specific range and distribution of protein binding site interactions. In addition, protein supersets optimized for independent compound libraries were cross-tested and were able to reproduce the performance of using the full protein library. This indicates that overtraining on a specific compound library during this iterative procedure was limited in scope, making these protein supersets broadly applicable for characterizing drug behavior.

## Methods

### Compound and indication mappings

The putative drug library in the CANDO platform comprises 3,733 human use compounds, including all clinically approved drugs from the US FDA, Europe, Canada, and Japan, collected from DrugBank [25], NCGC Pharmaceutical Collection (NPC) [26], Wikipedia, and PubChem [27]. Each drug/compound was converted to a three dimensional (3D) or tertiary structure to standardize input conformation and avoid biasing the results using ChemAxon’s MarvinBeans molconverter v.5.11.3 (https://chemaxon.com/). InChiKeys were generated from all preprocessed compounds using Xemistry’s Cactvs Chemoinformatics Toolkit [28] (https://www.xemistry.com/) to remove redundancies. Drug-disease associations were obtained from the Comparative Toxicogenomics Database (CTD) [29] and mapped to our drug/compound library, resulting in associations to 2,030 indications, including 1,439 indications with at least two drugs that are used to perform our leave-one-out benchmarking protocol described below [9–11, 13, 15].

### Compound-proteome interaction matrix generation

The protein structure library in the CANDO platform is made up of 48,278 tertiary conformations solved using x-ray diffraction taken from the Protein Data Bank (PDB) [30] (≈ 80% of the structures) as well as homology models (≈ 20%). The organism sources for the proteins include *Homo sapiens*, and several higher order eukaryotes, bacteria and viruses. Protein structure models were generated using HHBLITS [31], I-TASSER [32, 33], and KoBaMIN [34]. The I-TASSER modeling pipeline consists of the following steps: 1) HHBLITS and LOMETS for template model selection; 2) threading of protein sequences from templates as structural fragments; 3) replica-exchange Monte Carlo simulations for fragment assembly; 4) SPICKER [35] for clustering of simulation decoys; 5) ModRefiner [33] for generation of atomically refined model SPICKER centroids; 6) KobaMIN for final refinement of models. Some pathogen proteins failed during the modeling and were removed, ultimately resulting in 46,784 proteins in the final matrix. To generate scores for each compound-protein interaction, COFACTOR [36] was first used to determine potential ligand binding sites for each protein by scanning a library of experimentally determined template binding sites with the bound ligand from the PDB. COFACTOR outputs multiple binding site predictions, each with an associated binding site score. For each predicted binding site, the associated co-crystallized ligand is compared to each compound in our set using the OpenBabel FP4 fingerprinting method [37], resulting in a structural similarity score. The score that populates each cell in the compound-protein interaction matrix is the maximum value of all of the possible binding site scores times the structural similarity scores of the associated ligand and the compound.

### Benchmarking protocol and evaluation metrics

The compound-compound similarity matrix is generated using the root mean square deviation (RMSD) calculated between every pair of compound interaction signatures (the vector of 46,784 real-value interaction scores between a given compound and every protein in the library). Two compounds with a low RMSD value are hypothesized to have similar behavior [9–11, 13, 15]. For each of the 1,439 indications with two or more associated drugs, the leave-one-out benchmark assesses accuracies based on whether another drug associated with the same indication can be captured within a certain cutoff of the ranked compound similarity list of the left-out drug. This study primarily focused on a cutoff of the ten most similar compounds (“top10”), the most stringent cutoff used in previous publications [9–11, 13, 15]. The benchmarking protocol calculates three metrics to evaluate performance: average indication accuracy, compound-indication pairwise accuracy, and coverage. Average indication accuracy is calculated by averaging the accuracies for all 1439 indications using the formula c/d × 100, where c is the number of times at least one drug was captured within the cutoff (top10 in this study) and d is the number of drugs approved for that given indication. Pairwise accuracy is the weighted average of the per indication accuracies based on how many drugs are approved for a given indication. Coverage is the count of the number of indications with non-zero accuracies within the top10 cutoff.

### Superset creation and benchmarking

The 46,784 proteins in the CANDO platform were randomly split into 5,848 subsets of 8 and subsequently benchmarked using the method described above. The size of 8 was selected because offered the widest range of benchmarking values (relative to larger sizes), reduced the computational cost of the experiments (relative to smaller sizes that increase the number of individual benchmarks that need to be evaluated), divided into 46,784 evenly, and also provided adequate signal for the multitargeting approach to work according to our prior studies [12]. A total of 50 iterations were performed that resulted in 292,400 benchmarking experiments. Each subset was then ranked according to top10 average indication accuracy, pairwise accuracy, and coverage. The fifty best performing subsets from each ranking criterion (average indication accuracy, pairwise accuracy, and coverage) were progressively combined into supersets and benchmarked after each of the 50 iterations of the splitting and ranking protocol. The subsets were nonredundantly combined such that if a given protein was represented in the best performing subsets more than once (from two or more different iterations), then it would only occur once in the superset. The number of proteins in each superset ranged from 8 to 400.

### Independent compound library creation and evaluation

The CANDO putative drug library was split into two disjoint libraries comprising ≈50% of the compounds (1,866 and 1,867 compounds) five times. For each such library, the indication mapping was reconstructed to include only the disease associations of the drugs present. Each corresponding pipeline comprised of a disjoint compound library was subjected to ten iterations of the splitting and ranking protocol. Protein supersets, composed of the fifty best performing subsets for each metric that were progressively combined (see previous section), were generated and benchmarked. Supersets less than size 80 were excluded due to the results of Fig 2, which shows that benchmarking performance begins to plateau starting with around that number of proteins. The remaining supersets were then cross-tested on their complementary sets to assess the generalizability of our selection and optimization protocol by comparing it to the corresponding control value based on using the full protein structure library (but with the same disjoint compound library). The benchmarking for each of these 50% disjoint compound libraries are not directly comparable to each other nor to the full compound library because the indication mappings are different.

### Evaluating the features responsible for protein subset and superset accuracy

The best performing protein subsets and supersets were further analyzed to elucidate the protein feature(s) responsible for increased benchmarking performance. The protein subsets and supersets were analyzed based on four criteria: organismal source, structure source (x-ray diffraction or modeling), fold space (based on the CATH classification of proteins [38]), and interacting ligand structure distributions. The subsets and supersets were analyzed by counting the specific organisms to which the proteins belonged to see if any were over or underrepresented in the best and worst performing sets. Similarly, the subsets and supersets were analyzed to see if structures obtained via a specific source, x-ray diffraction or modeling, were differentially represented in the best and worst performing sets. Fold assignments were made to each protein in the subsets and supersets which were again analyzed for differential representation of specific protein folds. Finally, since our compound-protein interaction scoring method utilized the structural similarity of each drug/compound to the ligand co-crystallized with the protein (see previous section), we analyzed these ligands for differential representation. Each co-crystallized ligand in the COFACTOR database of template binding sites were clustered at various distances (0.1 to 0.9 with increments of 0.1) using the Butina clustering algorithm [39] in the RDKit [40] library based on the Tanimoto distance [41] of their Daylight fingerprints (http://www.daylight.com/). A total of 64,592 ligands were clustered with 7,252 ligands failing due to molecular file conversion errors. Each protein in the subsets and supersets was assigned a ligand cluster signature where each value in the vector is the number of times a ligand from a given cluster was chosen while calculating the compound-protein interaction score protocol for that protein. The fraction of interactions belonging to each cluster for each protein set was calculated by first adding the ligand cluster signatures for each protein belonging to the set, then ranking the clusters from greatest to smallest, and finally dividing by the number of total interactions (size of the protein set times 3,733). The ligand cluster signatures of the best and worst 50 subsets from the first iteration of the splitting and ranking protocol were averaged together, with the resulting distribution of cluster counts at each rank compared using Welch’s T-test for equal means. Protein subsets were chosen from only one iteration to ensure independence.

### Creating protein libraries using the most important feature

Of the features evaluated (see above), only the ligand cluster distribution could be correlated with benchmarking performance (see Results). We then generated new protein structure libraries that captured this feature ideally and assessed benchmarking performance. Each protein was ranked based on the variance of the values in their ligand cluster signatures. A minimum cutoff of two clusters was used to prevent the trivial case of proteins with only one cluster mapped (with a variance of zero). Another cutoff of at least 1,867 total mapped interactions was used to account for proteins with interactions mapped to unclustered ligands. Proteins were randomly selected from the top of the ranked list of variances at size increments of 20 and then benchmarked. The cutoff of proteins considered for random selection was two times the size of the library (for instance, 100 proteins were randomly selected from the top 200 proteins ranked by ligand cluster signature variance). A total of 50 libraries were made for each size to generate a distribution of benchmarking values. This procedure was repeated for the bottom of the ranked list of variances for comparison.

## Results

### Benchmarking of generated supersets

Fifty iterations of splitting the 46,784 proteins into 5,848 subsets of 8 were performed, resulting in 292,400 benchmarks. The maximum, minimum, mean, and standard deviation were respectively 11.7%, 6.2%, 9.1%, and 0.5% for average indication accuracy, 20.2%, 12.7%, 16.7%, and 0.8% for pairwise accuracy, and 548, 398, 477.5, and 15.6 for coverage with each metric following a normal distribution. The mean benchmarking performance for each superset within a given iteration tends to gradually increase as the number of iterations increases (Fig 1). Ranking subsets using coverage is overall the best criterion as it yielded the maximum benchmarking performance for all three metrics. Coverage was the dominant ranking criterion beginning at iteration sixteen as it yielded the maximum average indication accuracy and coverage. Average indication accuracy is overall the second best ranking criterion as evidenced by its very close performance to coverage in the average indication accuracy and pairwise accuracy metrics and being the best metric through iteration fifteen. The maximum values obtained by supersets for average indication accuracy, pairwise accuracy, and coverage, were 14.0%, 23.2%, and 602, which represents a 14%, 7%, and 10% improvement on the control (using all 46784 proteins) values of 12.3%, 21.6%, and 546, respectively.

**Fig 1.**
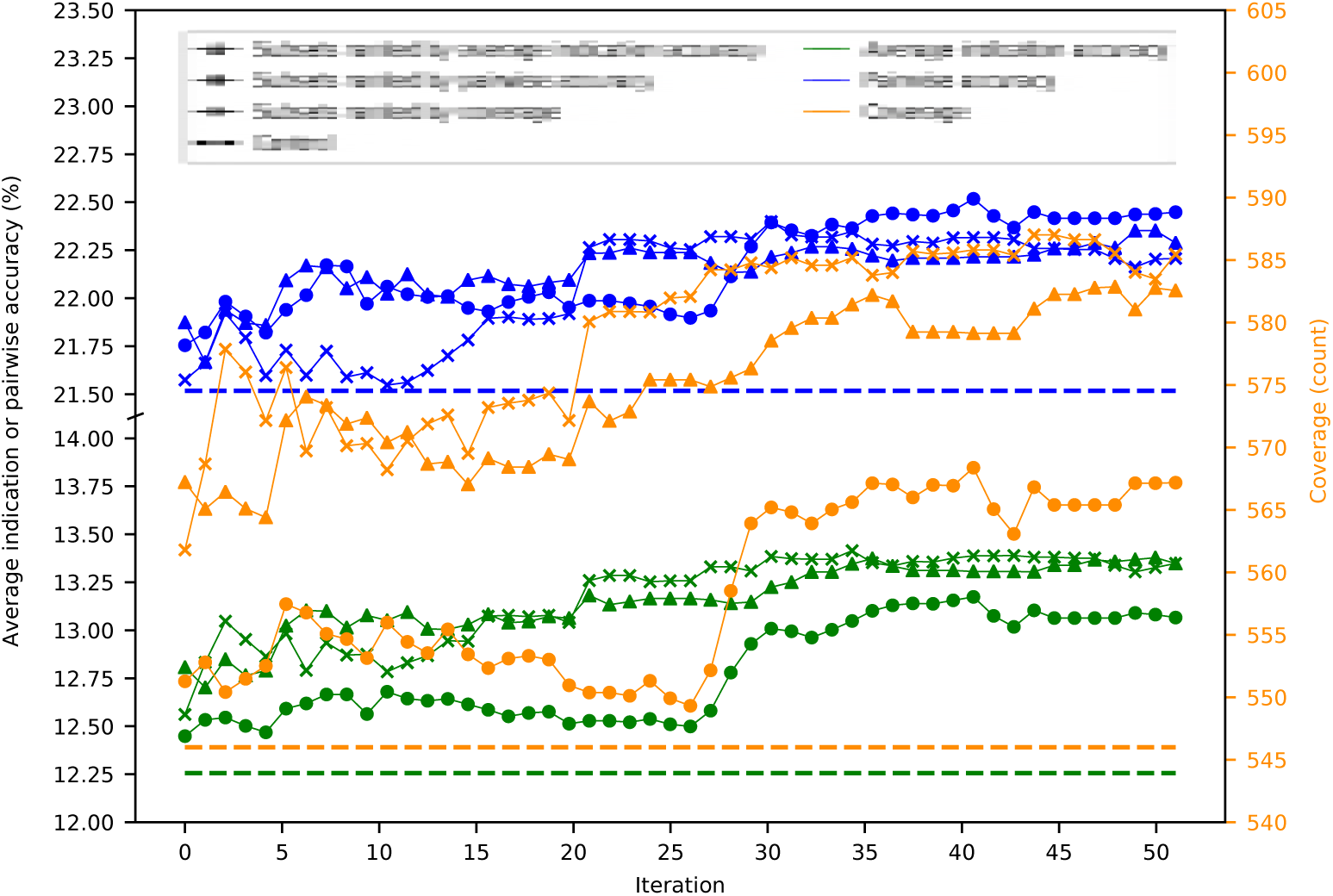
Mean superset benchmarking performance across 50 iterations. Each point represents the mean average indication accuracy (green), pairwise accuracy (blue), or coverage (orange) of 50 supersets created by consecutively combining the top 50 subsets ranked by average indication accuracy (triangles), pairwise accuracy (circles), or coverage (crosses). Average indication accuracy and pairwise accuracy are plotted on the left axis; coverage on the right. The dashed lines represent the control values for their respective metrics. The supersets improve on the control values and gradually increase in performance with the number of iterations with average indication accuracy being the best ranking criterion through ten iterations, after which coverage is superior, especially when measuring the coverage metric. This result demonstrates that the splitting and ranking protocol can produce supersets with benchmarking performance superior to using the full protein library by combining the best performing subsets with a vastly reduced number of proteins (100 to 1,000 fold reduction in size), further suggesting that specific groups of proteins are relatively more useful for drug repurposing accuracy.

**Fig 2.**
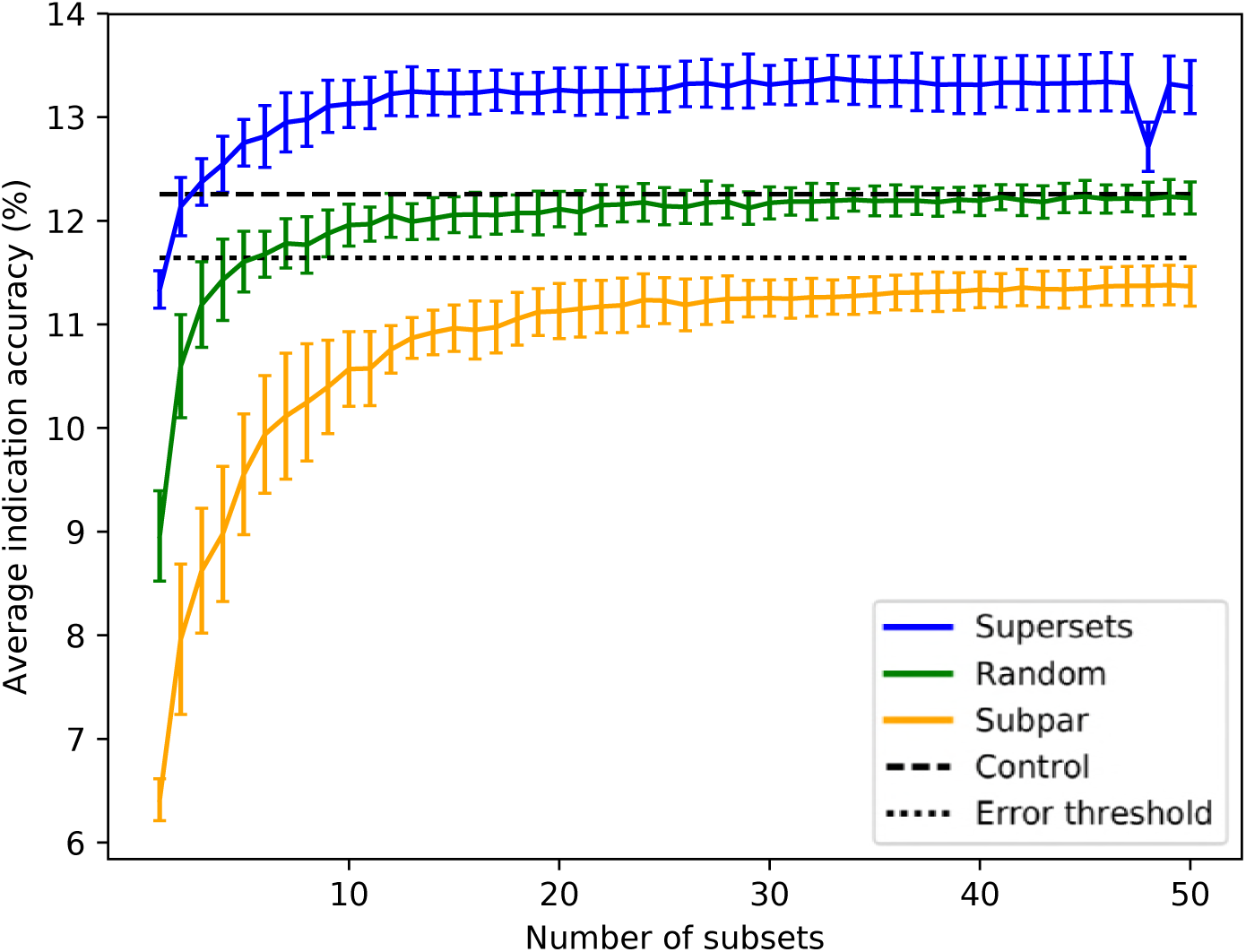
Superset, subpar set, and random set average indication accuracies sorted by size. Average indication accuracies are shown for the supersets (blue) generated using the best subsets ranked by the same metric. The line traces the mean score for each size with the bars indicating one standard deviation for the distribution. Subpar sets (orange) are the combinations of the worst performing subsets ranked by average indication accuracy. Randomly selected protein sets (green) of each size were also generated and benchmarked. The control value based on using the full protein library (dashed black at 12.3%) and an acceptable 5% threshold (dotted black at 11.6%) are plotted for reference (i.e., any protein set that benchmarks within 95% of the control value is considered acceptable). For the random sets and supersets, the performance in terms of average indication accuracy begins to plateau around 80-120 proteins. The supersets begin to slightly decline in performance after 32-33 subsets (256-264 proteins). The mean subpar set accuracies at each size all fall outside the 5% acceptable threshold, while the superset distributions are well above the control value with as few as five subsets. The difference between the superset and subpar set performance suggests that there is a particular distribution of features within their proteins that is correlated with benchmarking performance.

Sorting the supersets by size reveals that at least 80-120 proteins are required to reach optimal benchmarking performance (Fig 2). Nonredundantly combining the worst performing subsets of 8 into subpar sets demonstrates worse performance than the control value for the average indication accuracy based on using the full library, with the mean values of these subpar set distributions being below the acceptable 5% threshold of 11.6% for all sizes benchmarked. The random set and subpar set distributions begin to converge toward the average indication accuracy control value as size increases (Fig 2). The mean average indication accuracies begin to plateau or slightly decrease after size 256-264 for the supersets, while continuing to rise as the subpar set size increases. This suggests that the distribution of features within each protein library is important for describing drug/compound behavior.

### Cross-testing with independent compound libraries

For the independent compound library experiment, average indication accuracy was chosen as the ranking criterion because it performed the best in all three metrics through ten iterations in the superset experiment (Fig 1). Box and whisker plots for each compound library show the spread of the benchmarking performance for each metric generated from the protein supersets obtained through the splitting and ranking protocol of their complementary compound library (Fig 3). Supersets less than size 80 were excluded because there is a minimum number of protein features required to reach optimal benchmarking performance (Fig 2). The control value for the given metric fell within the inter-quartile range or below in 26 cases (87%), while it was within the upper quartile in the remaining four cases (13%), indicating that these values always fell within the range of the accuracy/coverage distributions. The control value was an extreme outlier below the distribution in five cases (17%), indicating the potential of these supersets to describe compound behavior more effectively than the full protein library.

**Fig 3.**
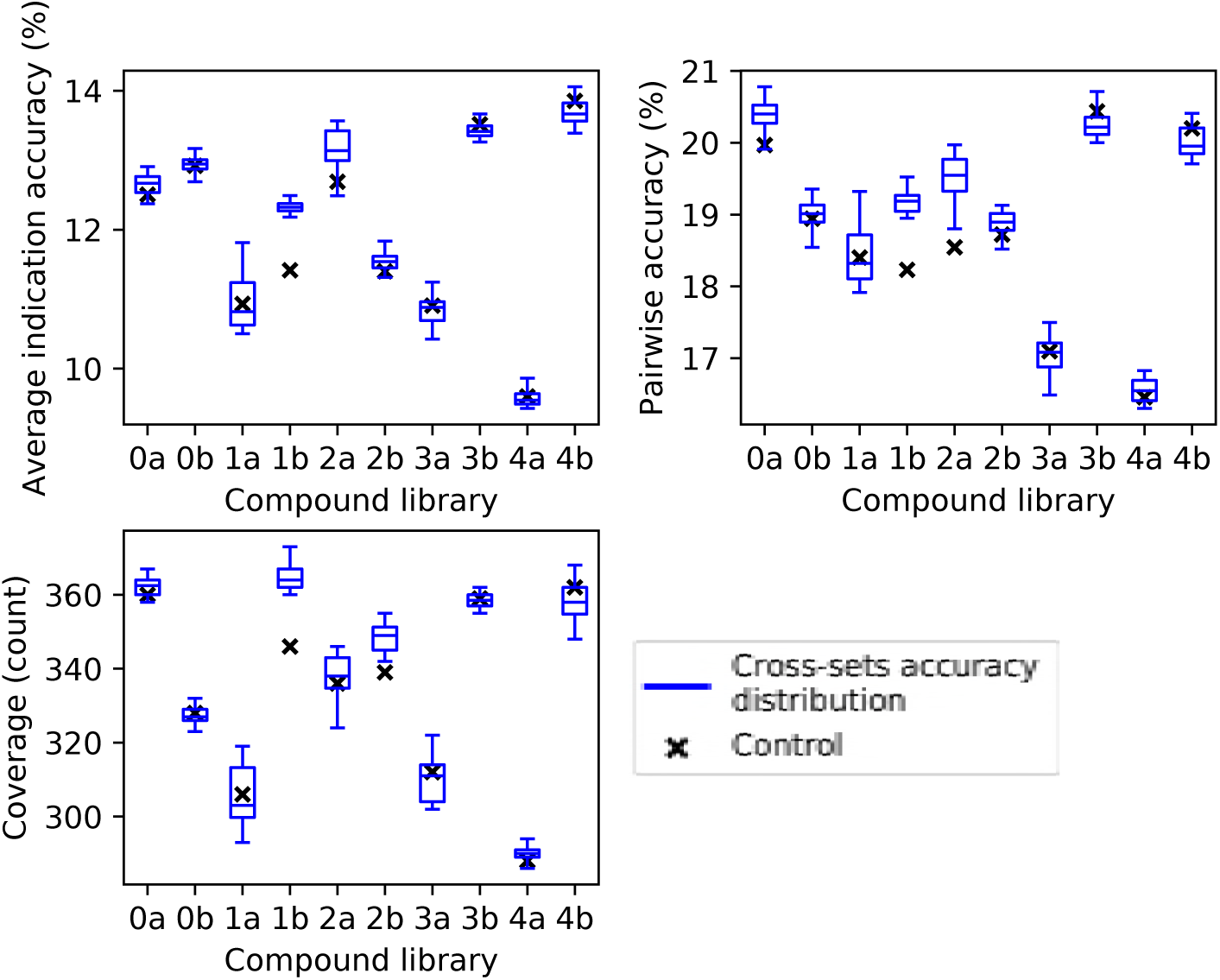
Benchmarking performance of protein supersets cross-tested with independent compound libraries. Protein supersets were generated using the splitting and ranking protocol with the average indication accuracy metric. Supersets were tested on their complementary library only (for instance, supersets generated from compound library 0a were tested on compound library 0b and vice versa). The blue box and whisker plots describe the benchmarking performance distributions of the 41 complementary supersets with the middle line being the median value, the box encompassing the first and third quartiles, and the whiskers extending to the maximum and minimum excluding outliers. The nine supersets less than size 80 were excluded due to results in Fig 2. In all cases, the control value for each compound library (black cross) never lies above the distribution of cross superset accuracies/coverages, indicating that these supersets can be generalized to any library of compounds.

### Ligand clustering and feature-based creation of protein libraries

The protein subsets and supersets were analyzed based on four criteria to elucidate the feature(s) responsible for benchmarking performance: organismal source, structure source (x-ray diffraction or modeling), fold space, and interacting ligand structure distributions. There were no significant correlations found for the first three criteria; no organism(s) or fold(s) was consistently represented in the best performing sets, while the the structure sources were proportionally distributed among the best and worst performing subsets and supersets. All ligands in the COFACTOR database were clustered to investigate the fourth criterion (see Methods). A total of 7,252 ligands were unable to be clustered due to molecular file conversion errors, resulting in 64,592 clustered ligands. Clustering at a distance of 0.3 created 9,929 clusters with varying sizes. The number of clusters including at least 10, 100, and 1,000 compounds were 568, 52, and 7, respectively, including 5,280 singleton clusters. Compound-protein interactions for all 46,784 proteins were mapped to ligand clusters based on the co-crystallized ligand of the binding site that was chosen for each compound-protein pair and ligand cluster signatures were generated for each protein (see Methods). The ligand cluster signatures of the best and worst 50 subsets from the first iteration of the splitting and ranking protocol were averaged together, with the resulting distribution of cluster counts at each rank compared using Welch’s T-test (Fig 4). The subsets with the best performance are composed of a much more equitable distribution of interactions among clusters than those with the worst performance on average. The first two ranks show the greatest contrast between the subsets with p-values of 1.04×10^*-*8^ and 2.38× 0^*-*4^ respectively, with all but two of the 22 rank distributions tested being significantly different (p-value < 0.05). The ligand cluster analysis was repeated for a superset and a subpar set and both follow a similar pattern to the one previously observed, with the superset having a more equitable distribution of interactions among clusters and the subpar set being far more imbalanced.

**Fig 4.**
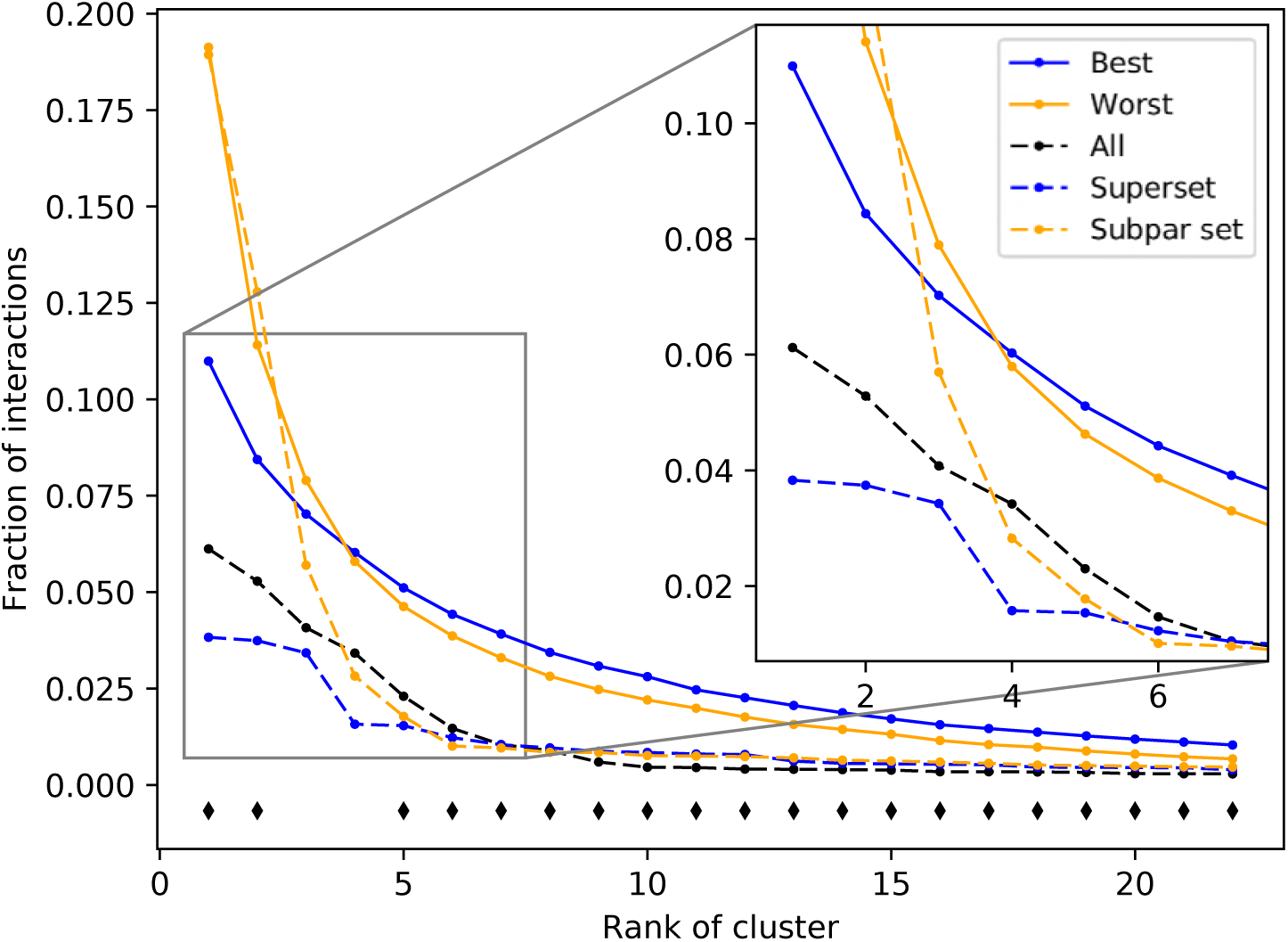
Fraction of interactions attributed to top ranked ligand clusters from different protein sets. The solid blue and solid orange points are averages of 50 best and worst subsets, respectively. Dashed lines represent an example of a superset (blue) and a subpar set (orange) which are nonredundant combinations of the best and worst performing subsets respectively. The control set (dashed black), representing the full protein structure library, falls in between the superset and subpar set. The black diamonds indicate that the distribution of counts at that cluster rank between the best and worst performing subsets, assessed using Welch’s T-test, is significant (p-value < 0.05). The subsets and supersets with the best performance demonstrate a more equitable distribution of interactions among ligand clusters as opposed to the worst performing subsets and subpar sets, indicating that using multitargeting proteins to compose our structure libraries yields superior benchmarking performance.

Based on the subset analysis, it was hypothesized that proteins having a more diverse and equitable distribution of interactions will benchmark better. Protein libraries were created by ranking the 46,784 proteins in CANDO on the variance of the number of interactions attributed to each ligand cluster (see Methods). Creating libraries using proteins with the lowest variance of ligand cluster interactions (excluding the trivial case of proteins mapped to only one cluster) results in benchmarking performance much higher than libraries composed of proteins with high variance (Figs 5 and 6). To elucidate the impact of the number of ligand clusters mapped to a given protein, we applied a upper cutoff filter for the number of ligand clusters allowed (Fig 5); all libraries were limited to size 80, which is the minimum required to reach optimal performance based on data in Fig 2. There is an optimal upper limit of ≈20-40 ligand clusters with regard to the benchmarking, indicating that using too many ligand clusters to describe a protein is undesirable for characterizing drug/compound behavior. Based on the data in Fig 5, we created protein libraries of various sizes using a upper cutoff of 40 (Fig 6). We begin to consistently recapture the benchmarking performance of the full library (within 5% error) with as few as 60 proteins. Libraries composed of high variance proteins, which are proteins with greatly imbalanced ligand cluster signatures (i.e. > 95% interactions mapped to one cluster), produce benchmarking performance outside of the acceptable 5% error range for all the created libraries (Figs 5 and 6). Fig 7 provides a visual depiction of the best and worst types of proteins for benchmarking performance, highlighting the idea that a moderately diverse and equitable distribution of interacting ligand clusters is ideal for describing drug behavior.

**Fig 5.**
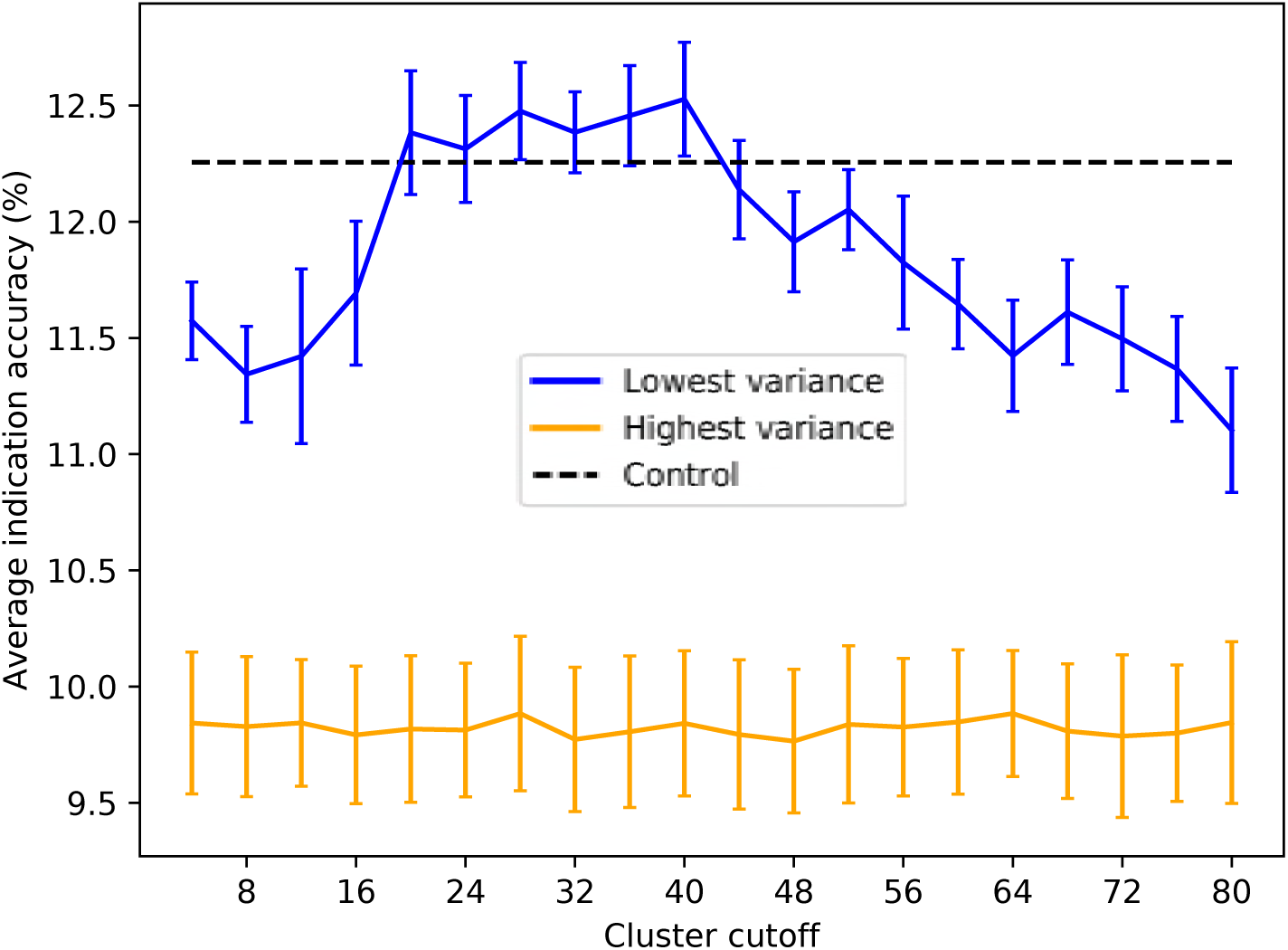
Average indication accuracy performance of created protein libraries of size 80 with various upper ligand cluster cutoffs. Protein libraries were created by randomly selecting 80 of the top 160 proteins from the list of proteins ranked by ligand cluster interaction variance (see Methods). A minimum of two ligand clusters was required to avoid the trivial case of only one cluster mapped (with a variance of zero). A upper ligand cluster count cutoff was applied to these protein libraries to determine the effect on benchmarking performance. The blue line traces the average indication accuracy distribution mean from benchmarking 50 libraries at each cutoff using the top ranked proteins. Similarly, the orange line traces the average indication accuracy distribution mean from benchmarking 50 libraries at each cutoff using the highest variance proteins. The bars indicate one standard deviation for the distribution. Using too small (< 20) or too large (> 40) of a cutoff results in suboptimal benchmarking performance. The high variance libraries result in average performance far below the acceptable 5% error range, with the cluster cutoff seemingly having no effect on average indication accuracy as all cutoffs produced comparable distributions. All average indication accuracies produced using a upper cutoff of 20-40 ligand clusters were within the acceptable 5% error range, with the upper cutoff of 40 ligand clusters producing the greatest mean accuracy. This result indicates that there is an optimal range of ligand cluster interactions to best describe therapeutic behavior.

**Fig 6.**
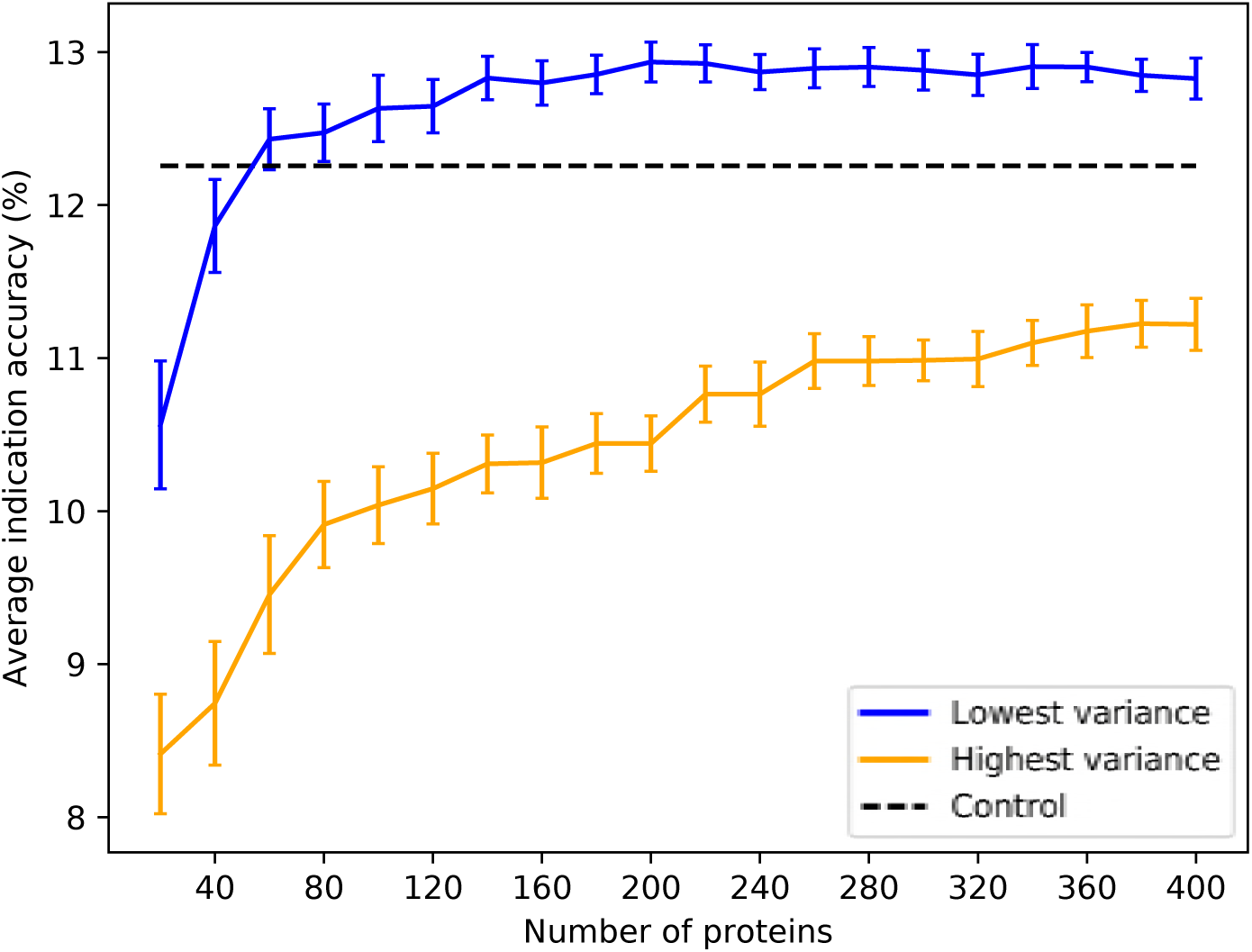
Average indication accuracy performance of created protein libraries at various sizes with ligand cluster cutoff of 40. Protein libraries were created by randomly selecting half the number of proteins in the library ranked by ligand cluster interaction variance with 2 to 40 ligand clusters mapped. The blue line traces the average indication accuracy distribution mean from benchmarking 50 libraries at each size using the top ranked proteins. Similarly, the orange line traces the average indication accuracy distribution mean from benchmarking 50 libraries at each size using the highest variance proteins. The bars indicate one standard deviation for the distribution. Using too small of a size (< 60) results in suboptimal benchmarking performance. Creating libraries from proteins with the highest variance results in performance on average far below the acceptable 5% error range, although size does have a positive correlation with performance with these high variance sets. This result reiterates that there is a minimum number of proteins required to reach optimal benchmarking performance and that proteins with high variance in their ligand cluster signatures are far superior for describing drug/compound behavior.

**Fig 7.**
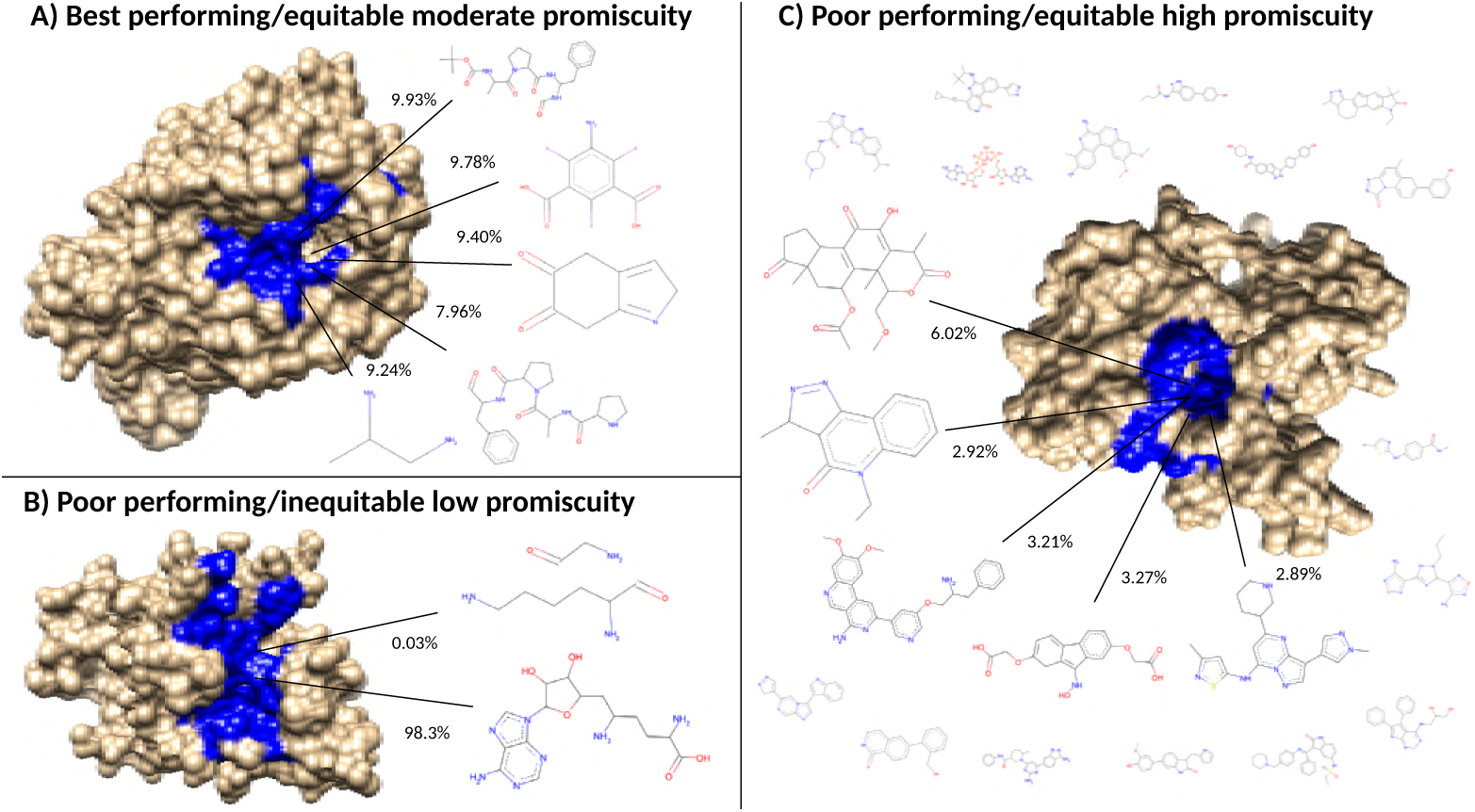
Visualization of the best and worst protein types for benchmarking performance based on ligand cluster signatures. Centroids of the top five ligand clusters from the signatures of each protein are depicted. The percent of interactions belonging to each ligand cluster are next to their respective centroid. Surface representations of the proteins were made using Chimera [42] with the predicted binding site residues from COFACTOR for each ligand shown (excluding the smaller ligands in C) colored in blue. A) Alkaline serine protease KP-43 from *Bacillus subtillus*: the top five ligand clusters account for 46.3% of the total interactions with the distribution between them being relatively equitable. B) SET domain of human histone-lysine N-methyltransferase: only two ligand clusters are predicted to interact, with one having over 98% of the total interactions. C) Human STE20-related kinase adapter protein beta: the ligand cluster signature is too promiscuous with the top five ligand clusters accounting for only 18.3% of the total interactions; the remaining sixteen ligands surrounding the protein account for 28.7%, which combined with the top five clusters is as much as the 46.3% of the total interactions shown in A from only five clusters. Subsets and supersets composed of proteins similar to A outperform those composed of proteins similar to B and C in benchmarking, indicating that moderately promiscuous proteins with equitable ligand cluster signatures are the best therapeutic descriptors.

## Discussion

The splitting and ranking protocol was originally intended to find a protein subset that benchmarked as least as well as the full set. The improvement of the benchmarking performance is an encouraging sign for incorporating machine learning in the CANDO platform in the future, and discovering how more complex weighting and relating of proteins contribute to drug repurposing accuracy, which is difficult to do with simple RMSD calculations. The smaller-sized protein libraries generated as part of this study, representing a 100 to 1,000 fold reduction in size, will be more conducive to machine learning. Feature reduction through the use of neural network based auto-encoders or principal component analysis will provide an important contrast to our proposed method.

The independent compound library experiment demonstrated that optimized protein sets based on a particular library were capable of therapeutically characterizing a completely different one, indicating that these supersets are generalizable. In other words, if a new drug/compound is added to the CANDO putative drug library, these reduced size supersets are likely able to describe its behavior at least as well as using every protein available. In addition to facilitating machine learning, our findings suggest a greatly reduced time required to generate new proteomic interaction vectors, which is particularly important if the program/protocol of choice for generating interactions is computationally expensive. Any repurposing candidates suggested from using the supersets are on average more clinically relevant as they were able to recapture drug behavior more accurately than using the full protein library in a statistically significant manner.

The knowledge that ligand cluster interaction diversity is the key requisite feature in describing drug behavior may also lead to optimization strategies other than the computationally intensive splitting and ranking protocol used in this study. The importance of ligand cluster interaction diversity may play a role in a variety of applications in systems and synthetic biology, for example in the design of protein systems that are specifically tailored to handle drug absorption, distribution, metabolism, and excretion processes. These optimizations incorporated into our repurposing platform should result in the rapid generation of more accurate putative drug repurposing candidates, thereby alleviating the problems associated with current drug discovery.

The ligand cluster analysis revealed that compound-protein interactions are more therapeutically relevant if the proteins used to describe the behavior of a compound are diverse in terms of the structures of the ligands which interact with their binding sites. Protein libraries with fewer predicted ligand cluster binding partners yield much worse performance than those consisting of proteins interacting with a more structurally diverse range of ligands. Coupling this with the finding that there is a minimum number of proteins required to reach optimal benchmarking accuracies (Fig 2), which was also observed by us previously [10, 11], drugs should realistically be described in the context of their multitarget nature, treating both small molecule compounds and proteins promiscuously, as in biological systems [43–45]. However, using libraries of proteins with too many diverse interactions in the CANDO platform also leads to suboptimal performance. We hypothesize this can be attributed to two factors: 1) spreading a compound interaction signature across too many (50 or more) ligand clusters can potentially dilute the therapeutic signal relative to the promiscuity of the corresponding proteins; or 2) these proteins are not therapeutically relevant and are therefore not useful for specifically describing drug behavior.

## Conclusions

We have developed an integrated pipeline that allows for the elucidation of proteins and their features that are important for benchmarking in the CANDO platform, and therefore important for drug repurposing. We are able to reproduce the performance of the complete CANDO protein structure library with orders of magnitude fewer proteins, allowing for more rapid candidate generation when evaluating new putative drug libraries or any other changes to the platform. We discovered that moderately promiscuous proteins, in terms of the structures of ligands with which they are predicted to interact, are important for describing how drugs behave in biological systems. Further optimization experiments using machine learning and indication-specific supersets are currently underway.

## Acknowledgments

The authors would like to acknowledge Dr. Zack Falls of the University at Buffalo for generating and processing much of the data used in this study.

